# A possible role for the kinetochore COMA complex in sensing the damage in *stn1-sz2*, a mutant of the Cdc13-Stn1-Ten1 telomeric complex

**DOI:** 10.64898/2026.10.02.756143

**Authors:** Nathalie Grandin, Michel Charbonneau

**Affiliations:** GReD Institute, CNRS UMR6293, INSERM U1103, Faculty of Medicine, University Clermont-Auvergne, 28 Place Henri Dunant, BP 38, 63001 Clermont-Ferrand Cedex, France

**Keywords:** budding yeast telomeres, Cdc13-Stn1-Ten1 complex, cell cycle checkpoints, mitotic tubulin spindle, kinetochore COMA

## Abstract

Telomeres, the ends of linear chromosomes, form a cap composed of specific telomeric DNA associated with specialized proteins that ensures efficient protection against chromosome degradation and, consequently, overall genome instability. Three main complexes have been implicated in chromosome end protection, namely telomerase, shelterin, and the CST complex (CTC1/Cdc13-STN1-TEN1). In a recent study, we uncovered mutants of the *S. cerevisiae* CST in which damage was sensed both by the two major spindle checkpoints and the major DNA damage checkpoint. In the best-studied of these mutants, *stn1-sz2*, the stability and organization of the mitotic tubulin spindle was profoundly affected, as were the organization of the centrosomes and the kinetochores. Here, we find that the *stn1-sz2* mutant is hypersensitive to deletion mutants of two members of the kinetochores, Ctf19 and Mcm21, which correlated with mis-localization of their associated subunit in the COMA complex, Okp1, during anaphase. Defects in interactions between the kinetochores and the tubulin spindle are entirely sensed by the SAC (spindle assembly checkpoint). Here, we find that some of the *stn1-sz2* damage is recognized by the Mad2 checkpoint protein independently from the SAC. The data presented here contribute to a better understanding of the damage in *stn1-sz2*, the prototype of novel mutants of telomeric CST characterized by mitotic-spindle-damage in addition to the classical telomeric-DNA-damage.

## 1. Introduction

Telomeres, the ends of linear chromosomes, recruit specialized proteins through binding to specific telomere DNA sequences present there. The main function of this protein-DNA telomeric complex is to form a cap that protects against chromosome degradation [1]. Indeed, in the absence of a functional telomeric cap, the telomeres would be recognized as DNA double-strand breaks by DNA damage-signaling and repair machineries. Chromosomes with unprotected telomeres undergo degradation, which leads to inappropriate recombination between the exposed sequences as well as to end-to-end fusion that cause chromosome breakage upon subsequent sister chromatid separation during the next mitosis. Another essential function of this telomeric cap is to recruit telomerase, a specialized reverse transcriptase enzyme that allows to replenish stocks of telomeric DNA sequences when needed. From the beginning of the studies on telomeres and telomerase up to now, the *Saccharomyces cerevisiae* budding yeast has been an excellent model organism, together with ciliated protozoans, *Drosophila melanogaster*, the *Schizosaccharomyces pombe* fission yeast, mouse and humans, principally [2].

In total, three protein complexes are needed to ensure telomere end protection in organisms possessing linear chromosomes, namely shelterin, CST and telomerase. Shelterin is a complex of six proteins, TRF1, TRF2, POT1, TIN2, TPP1, and RAP1, that prevents inappropriate recombination and fusion between telomeres [3-5]. Shelterin subunits also have complementary roles in telomere replication and length regulation. Shelterin is present in all vertebrates, as well as in the S. pombe fission yeast, but absent from the *S. cerevisiae* budding yeast. In *S. cerevisiae*, telomere end protection is principally taken in charge by CST [6,7]. The CST complex, initially thought to be present only in *S. cerevisiae*, was also identified in plants, mouse and humans, as well as in the *S. pombe* fission yeast [8,9] and, later, in the *Tetrahymena* ciliate [10] and the fruit fly *Drosophila* [11]. Finally, telomerase itself can be regarded as a telomere end protection complex, as it replenishes telomere sequences when needed, thus allowing the other two protection complexes to assume their function after having bound these sequences.

All three *S. cerevisiae* CST subunits are essential for telomere end protection and are also needed for telomere length regulation. Cdc13’s most important role is in telomerase recruitment to telomere ends, as well as in telomere end protection by capping. Stn1, in addition to its role in telomere end protection and telomere length regulation, also plays a specific role, being responsible, in association with DNA Polα, for the refilling of the strand previously elongated by telomerase [6]. Interestingly, Ten1, on the opposite, has no known telomeric specific function other than being essential for telomere protection as part of the CST complex. Most mutations in *S. cerevisiae* CST isolated to date caused extensive telomeric DNA degradation. The resulting accumulation of abnormally high levels of single-stranded telomeric DNA activated the Mec1/ATR DNA damage checkpoint, thus provoking a checkpoint-mediated cell cycle arrest, necessary for possible repair [12,13]. Recently, however, we isolated and started to characterize new *S. cerevisiae* mutants of CST that simultaneously exhibited defects that were sensed not only by the DNA damage checkpoint, but also by the two major spindle checkpoints, the so-called SAC (spindle assembly checkpoint) and SPOC (spindle position checkpoint) [14]. In one of these new CST mutants, *stn1*-*sz2*, the mitotic tubulin spindle was profoundly destabilized, with dramatic consequences on the organization of the spindle pole bodies (centrosomes) and kinetochores during mitosis [15]. Therefore, the CST complex might not only provide telomere end protection by masking the telomeric DNA from DNA processing enzymes, but also ensure full stability of the mitotic spindle during events occurring prior to the segregation of the sister chromatids in anaphase.

At this point of the characterization of these new mutants of CST, it is important to identify the sensors of the produced damage, as well as the targets of these sensors. In a candidate approach, we found, in the present study, that loss of function of two members of the kinetochores, Ctf19 and Mcm21, was strongly aggravating the damage generated by the *stn1*-*sz2* mutation. In the second part of the present study, we found that part of the *stn1*-*sz2* damage was recognized by Mad2, one of the major components of the SAC, and the Top2 checkpoint, but independent from its role in the SAC. Given the existing data from the literature, we propose that the so-called Top2 metaphase checkpoint might be activated by the *stn1*-*sz2* damage, in parallel with the SAC, both depending on Mad2. Altogether, the present data provide new information on the newly uncovered mitotic-spindle-damage functions of the chromosome end protection CST complex. These new functions of CST appear to be required later than the telomeric-DNA-damage functions, in anaphase rather than at the G2/metaphase stage.

## 2. Materials and methods

### 2.1. Yeast strains and media

The *Saccharomyces cerevisiae* yeast strains used in this study were derivatives of BF264-15Daub (*ade1 his2 leu2*-*3,112 trp1*-*1a ura3*Δ*ns*), described previously [16]. Yeast cultures were grown at the indicated temperatures in YEP (1% yeast extract, 2% bactopeptone, 0.005% adenine, and 0.005% uracil) supplemented with 2% glucose (YEPD) or in selective minimal media.

The GFP ORF (Open Reading Frame) sequences were attached in frame to the 3′ end of the *OKP1* gene cloned into one of the integrative expression vectors of the *S. cerevisiae* YIplac series [17]. Details for the construction of YIplac-*OKP1*-GFP and its integration at *OKP1* genomic locus will be available upon request.

The *SPC110*-td-Tomato and *tub2*-*430* strains were kindly provided by Jeffrey Moore. The *MAD2*-GFP strain was kindly provided by Duncan Clarke. The *stu1*-*5* strain was kindly provided by Tim Huffaker. The *bub2*Δ and *mad2*Δ mutants were kindly provided by Andrew Murray. The *bub1*Δ, *bub3*Δ, *ctf19*Δ, and *mcm21*Δ strains were purchased from Euroscarf (Oberursel, Germany). The *csm1* null strain was constructed by replacing the entire *CSM1* ORF by *URA3* (ORF + flanking sequences). Details of the construct can be made available upon request. All strains were made isogenic by back-crossing them at least five times against our genetic background.

Cell viability was determined by performing the so-called “drop tests” or “spot assays”. To perform these tests, cells from exponential growth cultures were counted with a hematocytometer, and the cultures were then serially diluted by 1/10^th^, spotted onto YEPD (or selective medium) plates, and incubated at the desired temperatures for 2 days before being photographed.

### 2.2. Fluorescence and confocal microscopy

Cells were observed using a Zeiss (Oberkochen, Germany) LSM 800 Airyscan confocal microscope [15]. Cells were mounted on slides in the Vectashield with DAPI mounting medium (Vector Laboratories, Inc; Newark, CA 94560, USA).

### 2.3. Statistical analyses

A Student’s t-test conducted using Prism software (version 7.0d) was used throughout the statistical analyses [15]. Significance numbers are given as p-values: ns stands for “nonsignificant”, * stands for a p-value < 0.05, ** stands for a p-value < 0.01, *** denotes a p-value < 0.001, and **** indicates a p-value < 0.0001.

## 3. Results

### 2.1 Implication of the Ctf19-Mcm21-Okp1 kinetochore subcomplex in spindle-associated Stn1 functions

Our recent study established that *stn1*-*sz2* mutant cells were enriched, in comparison with the *cdc13*-*1* mutant (the best studied CST mutant), in the proportion of cells in anaphase [15]. The kinetochore consists of a large multiprotein complex arranged around the centromeric structures of chromatids to attach to microtubules of the mitotic spindle until they can be correctly segregated at anaphase [18]. We previously observed that the kinetochores were abnormally attached to the spindle in the *stn1*-*sz2* mutant [15].

These observations led us to hypothesize that members of the kinetochore complex might play a role in detecting and/or transmitting the damage signals present in the *stn1*-*sz2* mutant. Interestingly, in a candidate approach, *stn1*-*sz2* mutant cells exhibited strong negative genetic interactions with mutants bearing a deletion of either the *CTF19* or *MCM21* genes, coding for the two non-essential subunits of the kinetochore COMA complex (**Figure 1A**). Thus, the presence of Ctf19 or of Mcm21, which is not required in otherwise wild-type cells, becomes essential in temperature-sensitive *stn1*-*sz2* mutant cells, even at their permissive temperatures for growth (**Figure 1A**). This strongly suggested that Ctf19 and Mcm21 intervene in molecular mechanisms that act in controlling some of the damage generated by the *stn1*-*sz2* mutation.

**Figure 1.**
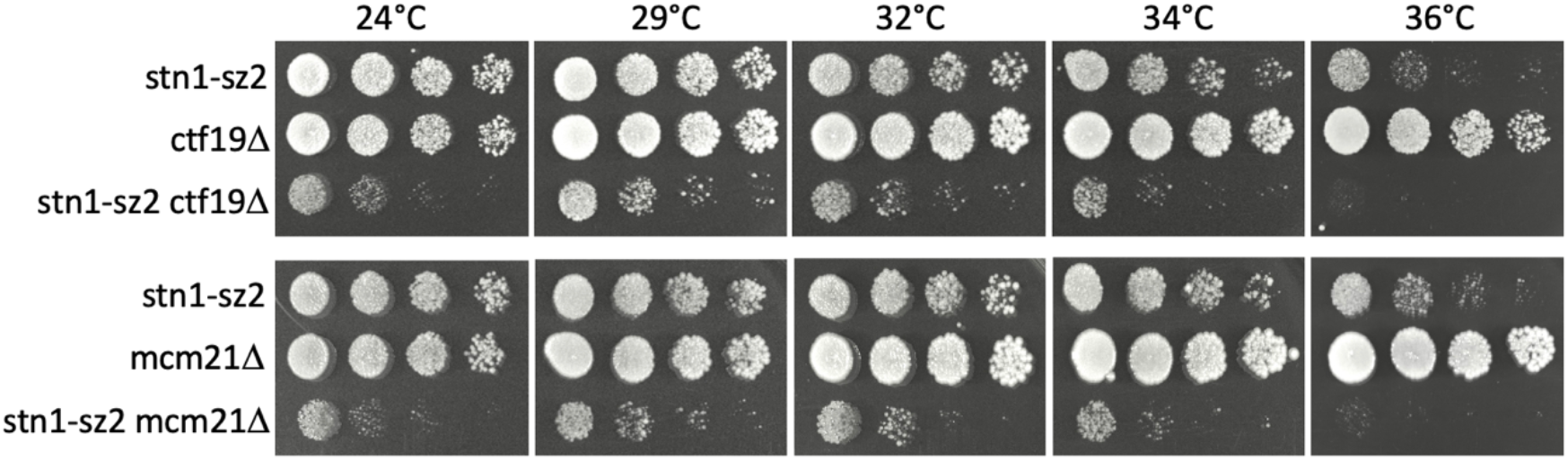
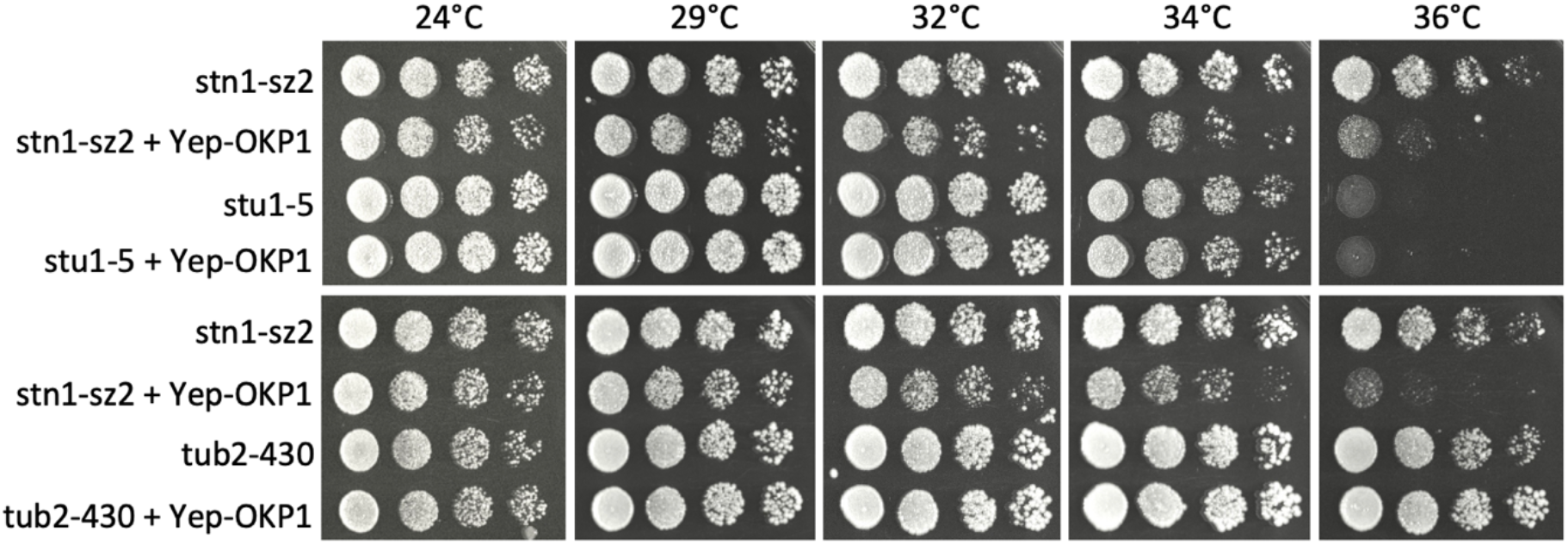
Genetic interactions between the temperature-sensitive *stn1*-*sz2* mutant and null (deletion) mutants of *CTF19* or *MCM21*. (**A**) The “synthetically lethal” effects of the *ctf19*Δ or *mcm21*Δ on the growth properties of the *stn1*-*sz2* mutant cells were well visible at all indicated temperatures, when comparing the extent of growth between the double mutant and that of each of the single mutants (see text for explanations). (**B**) Overexpression of *OKP1* (“Yep-OKP1”, from a yeast episomal plasmid) resulted in an aggravation of viability of the temperature-sensitive *stn1*-*sz2* mutant, an effect that started to be visible at 29°C (rows 1 and 2 of each, top and bottom, panels). Control experiments were performed using two additional temperature-sensitive mutants, *stu1*-*5* (top panel) and *tub2*-*430* (bottom panel), as indicated. Overexpression of *OKP1* was driven by a multicopy (episomal, 2μ, plasmid). “Spot assays” (or “drop tests”) were performed using ten-fold serial dilutions (from left to right) of yeast cultures grown for 2 days on YEPD plates at the indicated temperatures.

Although Ctf19 and Mcm21 are non-essential proteins, the other two components of COMA, Okp1 and Ame1, are indispensable for cell survival [19]. In experiments aiming to decipher the putative role of Ctf19 and Mcm21 in “handling” *stn1*-*sz2* damage, we found that overexpression of *OKP1* from a multi-copy plasmid clearly aggravated the growth properties of the *stn1*-*sz2* mutant cells (**Figure 1B**). Control experiments established that overexpression of *OKP1* did not alter the growth properties of temperature-sensitive mutants of spindle functions, such as *stu1*-*5* and *tub2*-*430* (**Figure 1B**).

Due to the great difficulty to explain the effects of manipulating the expression of *OKP1* on the growth properties of the *stn1*-*sz2* mutant, we turned to a somewhat more direct approach and set out to visualize Okp1 localization in the wild type versus the *stn1*-*sz2* mutant. By confocal microscopy, Okp1 was visualized using an *OKP1*-GFP construct, while, simultaneously, the spindle pole bodies (SPBs, the homologs of the centrosomes in higher eukaryotes) were visualized using an *SPC110*-td-tomato construct (**Figure 2**).

**Figure 2.**
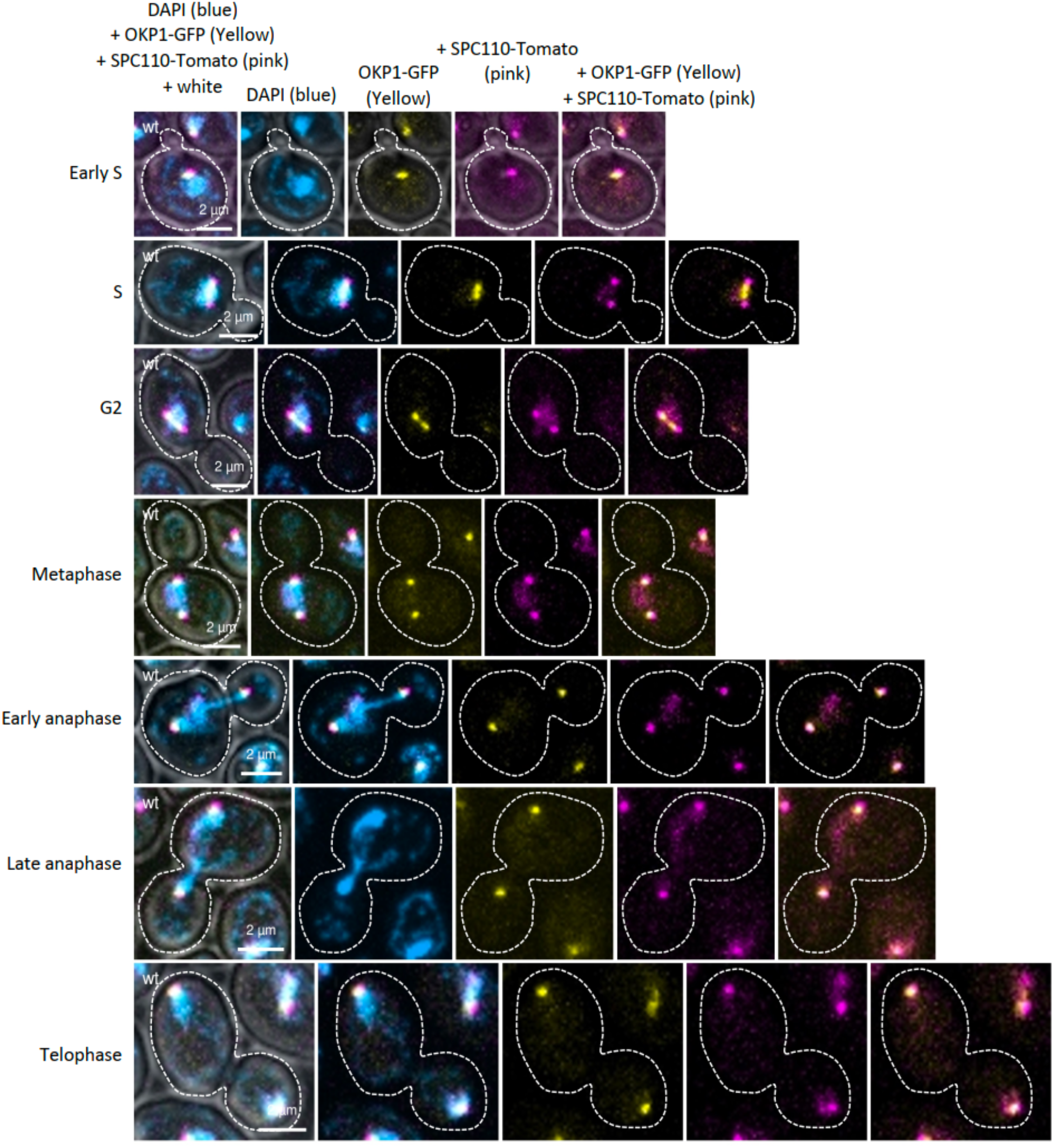
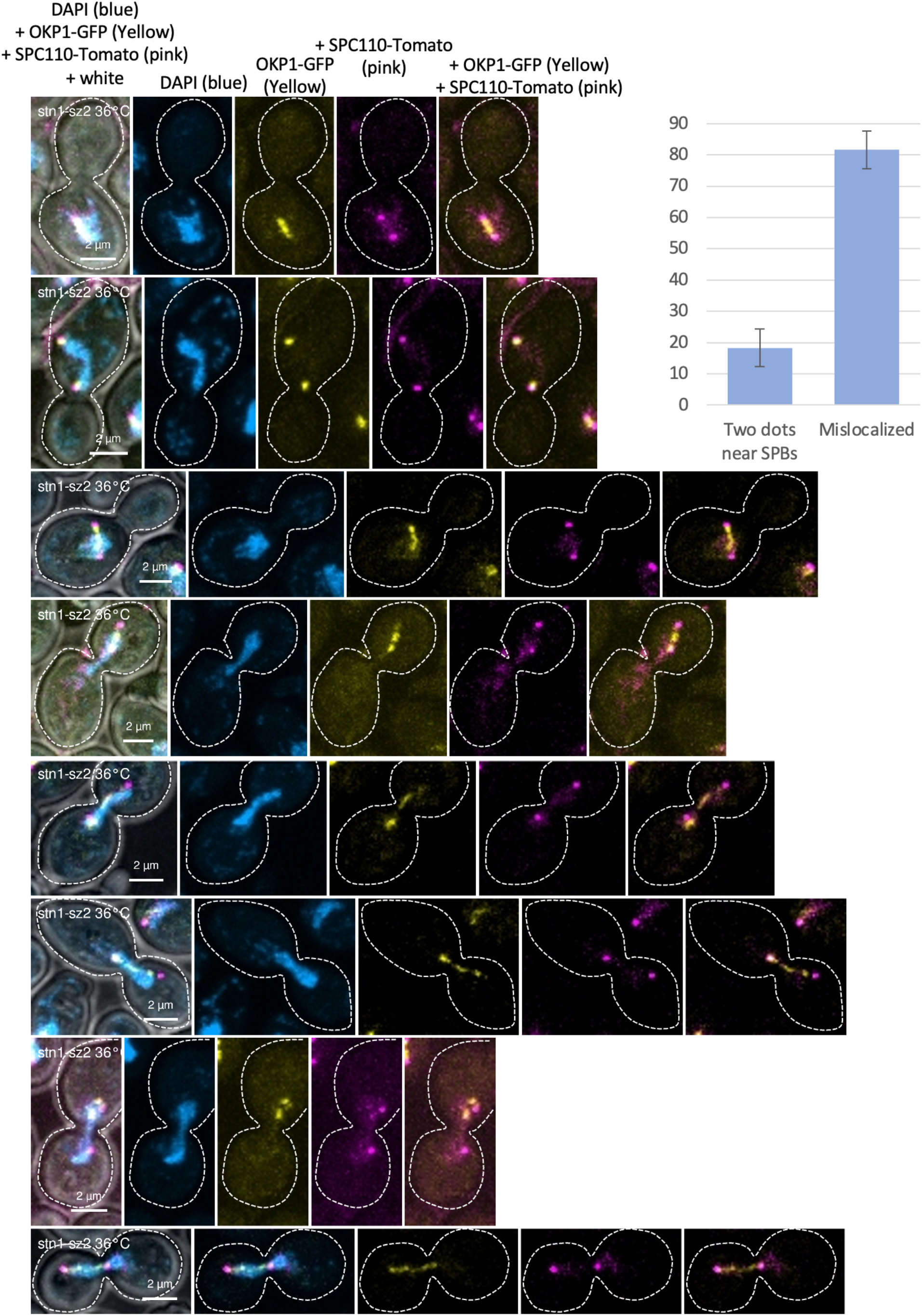
Localization of endogenous Okp1-GFP (shown with yellow labeling rather than green, for better visualization) throughout the cell cycle of wild-type cells (**A**) and in *stn1*-*sz2* mutant cells (**B**) at the fully restrictive temperature for growth of 36°C. All cells were also bearing an *SPC110*-Tomato integrated construct in order to visualize the spindle pole bodies (SPBs), the centrosomes homologs (represented in pink, rather than red, for better visualization). DAPI staining was used to simultaneously visualize nucleus DNA (shown with blue labeling). See main text for explanations. The histograms of panel B show the quantification of the mislocalized Okp1-GFP signals in *stn1*-*sz2* cells after 2 hr growth at 36°C. Results from 6 independent experiments are shown as percentages of total cells (100 cells counted per experiment).

In wild-type cells (**Figure 2A**), Okp1-GFP localized at the SPBs during early S phase, and then along the axis joining the SPBs (which corresponds to the mitotic spindle) when SPBs dissociated following their duplication. As mitosis progressed, the Okp1-GFP signal re-associated with kinetochores that were pulled to near the SPBs.

In *stn1*-*sz2* mutant cells grown for 2 hours at the restrictive temperature of 36°C, which arrested cell cycle progression in metaphase/early anaphase of the cell cycle, Okp1-GFP localized just like in the wild type in early metaphase, but no longer in anaphase (**Figure 2A, B**). Indeed, in *stn1*-*sz2* it remained along the spindle, contrary to the situation in the wild type in which it always remained near the SPBs, like the kinetochores do at this stage (**Figure 2**). Therefore, in *stn1*-*sz2*, the Okp1-bound kinetochores appeared not to be correctly pulled toward the SPBs. This observation confirmed our precedent finding that Slk19-GFP was often seen away from the spindle axis [15]. Slk19 binds to kinetochores not attached to the spindle, therefore away from the spindle axis [20]. Therefore, we have an indication that, in the *stn1*-*sz2* mutant, the kinetochores that were correctly attached to the spindle were restrained in mobility toward the SPBs.

### 3.2. Genetic interactions between STN1 and CSM1

Functional interactions between Stn1 and Csm1, on the one hand, and between Ctf19 and Csm1, on the other hand, have been previously described in the literature. Thus, in a genome-wide two-hybrid screen, Stn1 was found to physically interact with Csm1 [21]. In addition, Ctf19 was also shown, in the same screen, to bind Csm1 [21]. Interestingly, Csm1 was found to localize at the level of the rDNA sequences (inside the nucleolus) during metaphase and to relocate to the kinetochores during anaphase [22].

Given the data above concerning the implication of Ctf19 in the response to *stn1*-*sz2* damage, we reasoned that it would be interesting to know whether genetic interactions between *STN1* and *CSM1* might exist. This was indeed the case, since we found that the *stn1*-*sz2 csm1* disruption (*CSM1* is a non-essential gene) double mutant exhibited impaired viability compared with the two single mutants at all temperatures between 24 and 36°C (**Figure 3**). Therefore, the presence of Csm1, which is not required in otherwise wild-type cells, becomes essential in temperature-sensitive *stn1*-*sz2* mutant cells, even at their permissive temperatures for growth.

**Figure 3.**
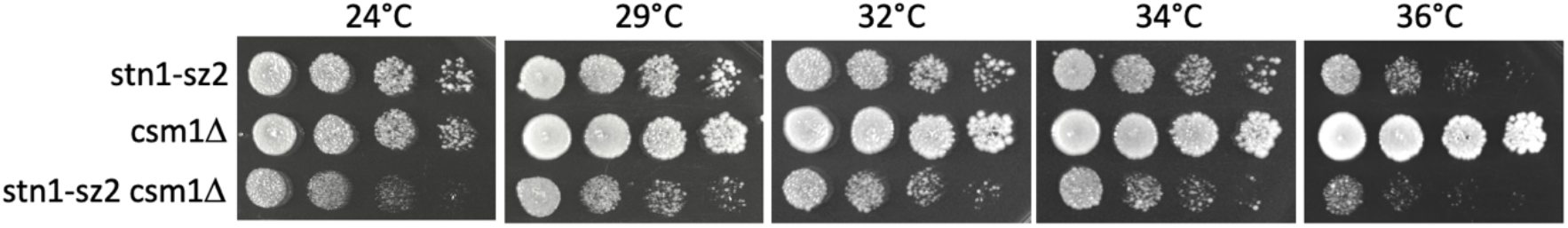
Genetic interactions between the temperature-sensitive *stn1*-*sz2* mutant and the null (deletion) mutant of *CSM1*. The “synthetically lethal” effects of the *csm1*Δ on the growth properties of the *stn1*-*sz2* mutant cells were well visible at all indicated temperatures, when comparing the extent of growth between the double mutant and that of each of the single mutants (see text for explanations).

### 3.3. In the stn1-sz2 mutant, Mad2 does not massively re-localize at kinetochores unlike in nocodazole-treated cells

The *stn1*-*sz2* mutant has the particularity to activate the two major mitotic spindle checkpoints, contrary to all previously identified mutants of CST that only activate the major G2/M DNA damage checkpoint [15]. The two major spindle checkpoints are the spindle position checkpoint (SPOC), whose central subunit is Bub2 (in complex with Bfa1), and the spindle assembly checkpoint (SAC), which principally implicates Mad1, Mad2, Mad3, Mps1, Bub1 and Bub3.

An interesting observation was that the effects of Mad2 or Mad1 inactivation in response to *stn1*-*sz2* damage in terms of cell viability was different from their response to microtubule-depolymerizing agents, benomyl or nocodazole. Thus, inactivation of *MAD2* or of *MAD1* in the *stn1*-*sz2* mutant resulted in an increase in cell viability, indicated by the improvement of colony formation, an effect best seen at 34-36°C (**Figure 4A**), as also shown before [14]. In contrast, in case of damage provoked by benomyl, the presence of the *mad2* or of the *mad1* deletion provoked a decrease in cell viability [23,24]. The phenotype of the *stn1*-*sz2 mad2*Δ and *stn1*-*sz2 mad1*Δ mutants were typical of that of mutants forcing the passage of checkpoint-mediated cell cycle arrest. This happens when the damage activating the checkpoint can be tolerated through the following mitosis. These events have been best described concerning mutants of *cdc13*-*1* defective in the G2/M DNA damage checkpoint [12,25]. On the other hand, the phenotype of benomyl-treated *mad2*Δ or *mad1*Δ mutants were more classical, exhibiting loss of viability provoked by the failure of these damaged cells to properly arrest cell cycle progression, due to inactivation of the SAC [23].

**Figure 4.**
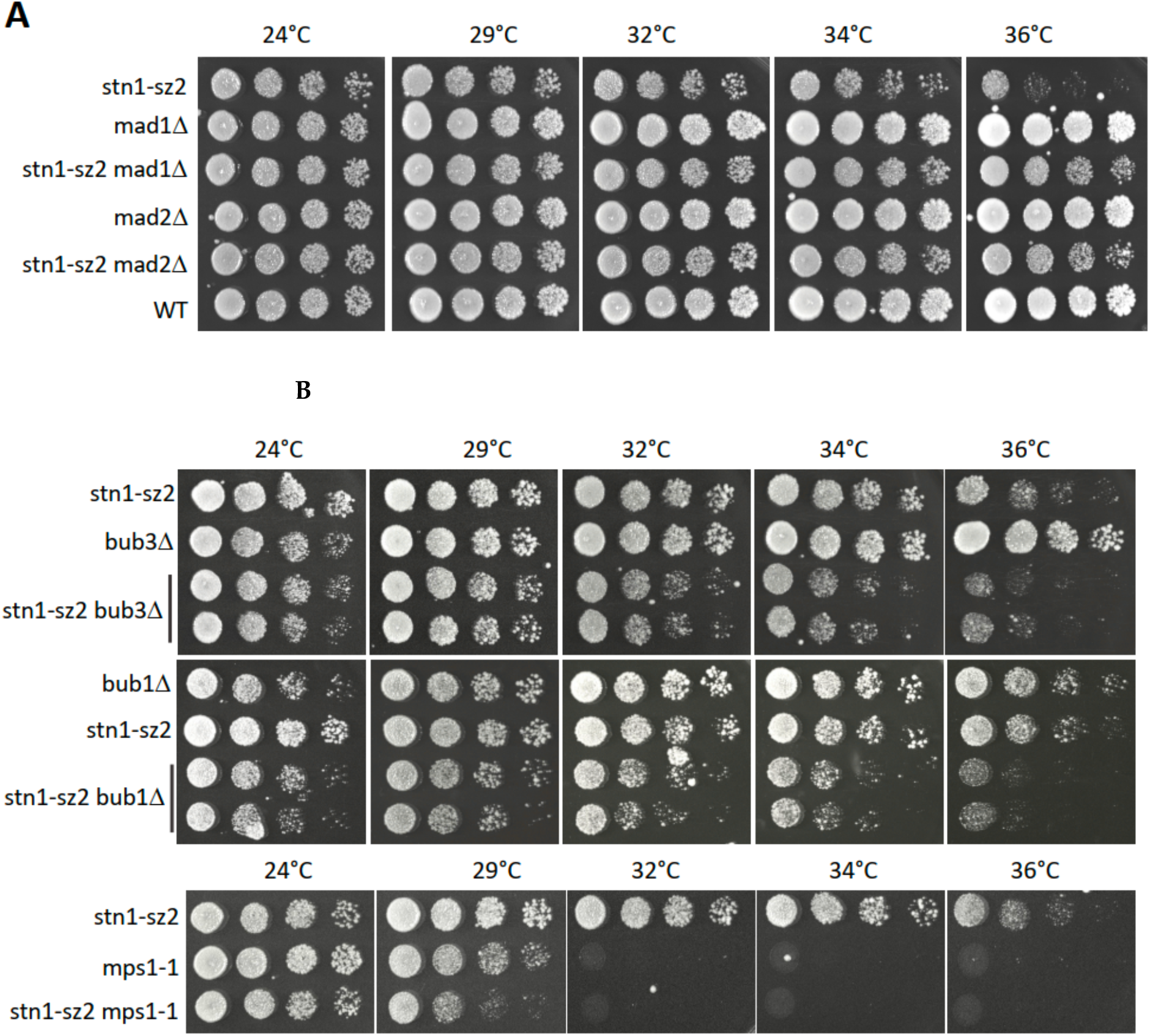
Detection of *stn1*-*sz2* defects by various components of the SAC (spindle assembly checkpoint) and of the SPOC (spindle position checkpoint) (**A**) Inactivation of the SAC in the *stn1*-*sz2* mutant, achieved in the presence of the mad1Δ or mad2Δ mutation background, resulted in an improvement of viability, as best seen here at 36°C. “WT” represents wild-type cells. (**B**) In contrast, genetic inactivation of components of the SAC other than Mad1 or Mad2, achieved in the *BUB1*, or *BUB3* or *MPS1* mutation background, resulted in an aggravation of the growth properties of the *stn1*-*sz2* mutant cells. Note that *mps1*-*1* is a temperature-sensitive mutant. See text for explanations. “Spot assays” (or “drop tests”) were performed using serial tenfold dilutions (shown from left to right) of liquid cultures grown for 2 days on YEPD plates at the indicated temperatures.

In the response to spindle damage, classical studies have placed Bub1, Bub3 and Mps1 upstream of Mad1 and Mad2 in the SAC pathway [26]. Interestingly, the *stn1*-*sz2 bub1*Δ, *stn1*-*sz2 bub3*Δ and *stn1*-*sz2 mps1*-*1* mutants all exhibited a classical checkpoint phenotype, that is to say aggravated cell mortality of the double mutants when compared with the two corresponding single mutants, a phenomenon known as synthetic lethality or synthetic sickness (**Figure 4B**). In addition, inactivation of the Bub2 SPOC component in *stn1*-*sz2* also provoked decreased cell viability [14], similar to the effect of the *bub2* deletion on benomyl-treated cells [27]. Again, this was provoked by the failure of these damaged cells to properly arrest cell cycle progression, due to inactivation of the SPOC [27].

In summary, the *mad2*Δ and *mad1*Δ mutations elicit a completely different response in *stn1*-*sz2* mutant cells versus in cells treated with benomyl or nocodazole. On the other hand, inactivation of all other SAC subunits Bub1, Bub3 and Mps1, as well as of the Bub2 SPOC subunit had the same effect in the *stn1*-*sz2* mutant and in benomyl-treated cells.

The increased growth properties and viability provoked by the *mad2*w or *mad1*w mutation in the *stn1*-*sz2* mutant have, to our knowledge, never been observed in other instances of spindle damage. We hypothesized that the *stn1*-*sz2* damage triggered an additional mitotic checkpoint distinct from the SAC, but also implicating Mad1-Mad2. To test this hypothesis, we first set out to determine the localization of Mad2-GFP in *stn1*-*sz2* mutant cells and in nocodazole-treated cells, both in comparison with that in untreated wild-type cells (**Figure 5**). Mad1 and Mad2 are effectors of the SAC that localize to the kinetochores only when the SAC is activated. In the absence of SAC-recognized damage, they are located at the nuclear envelope [28,29].

**Figure 5.**
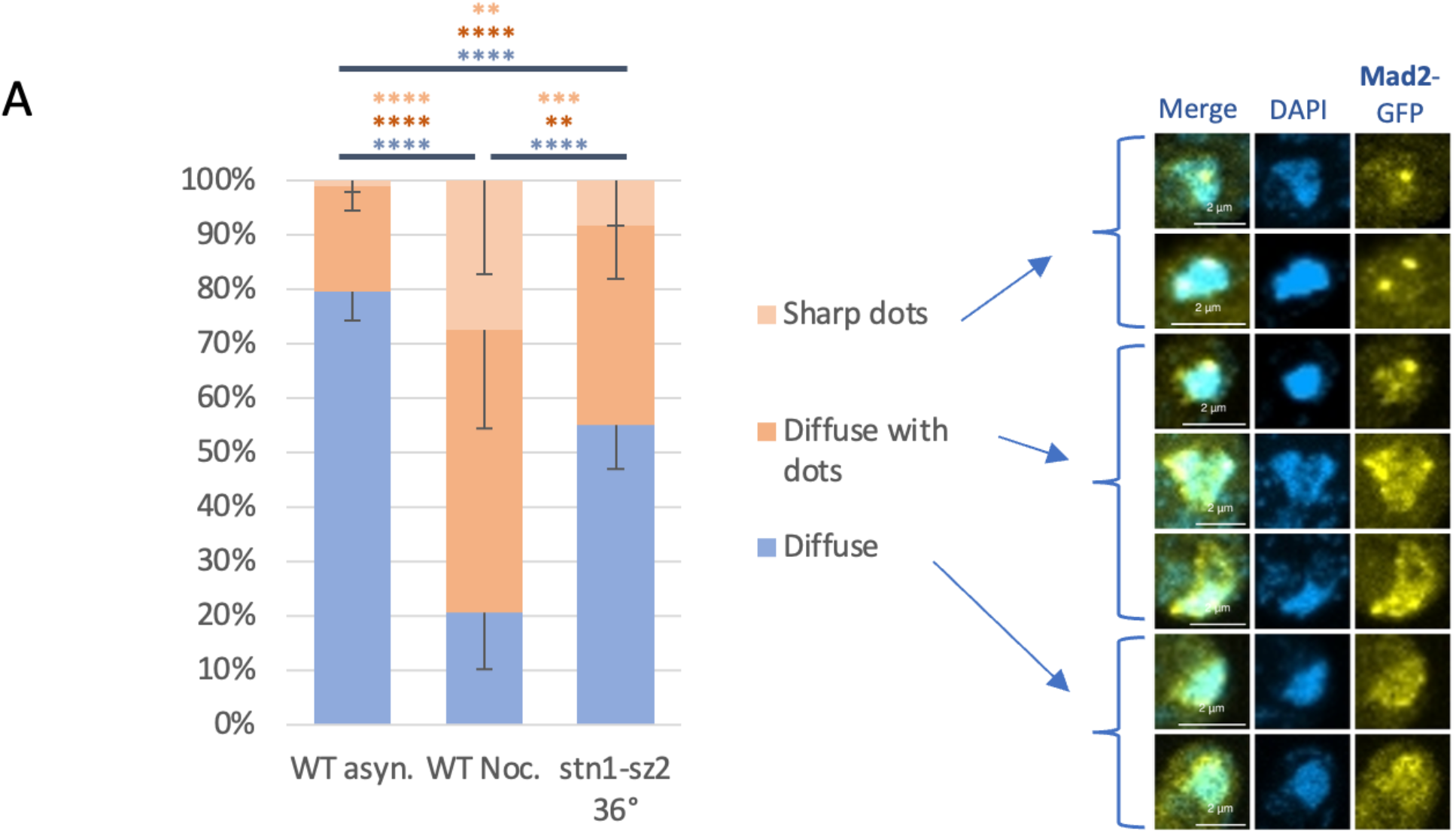
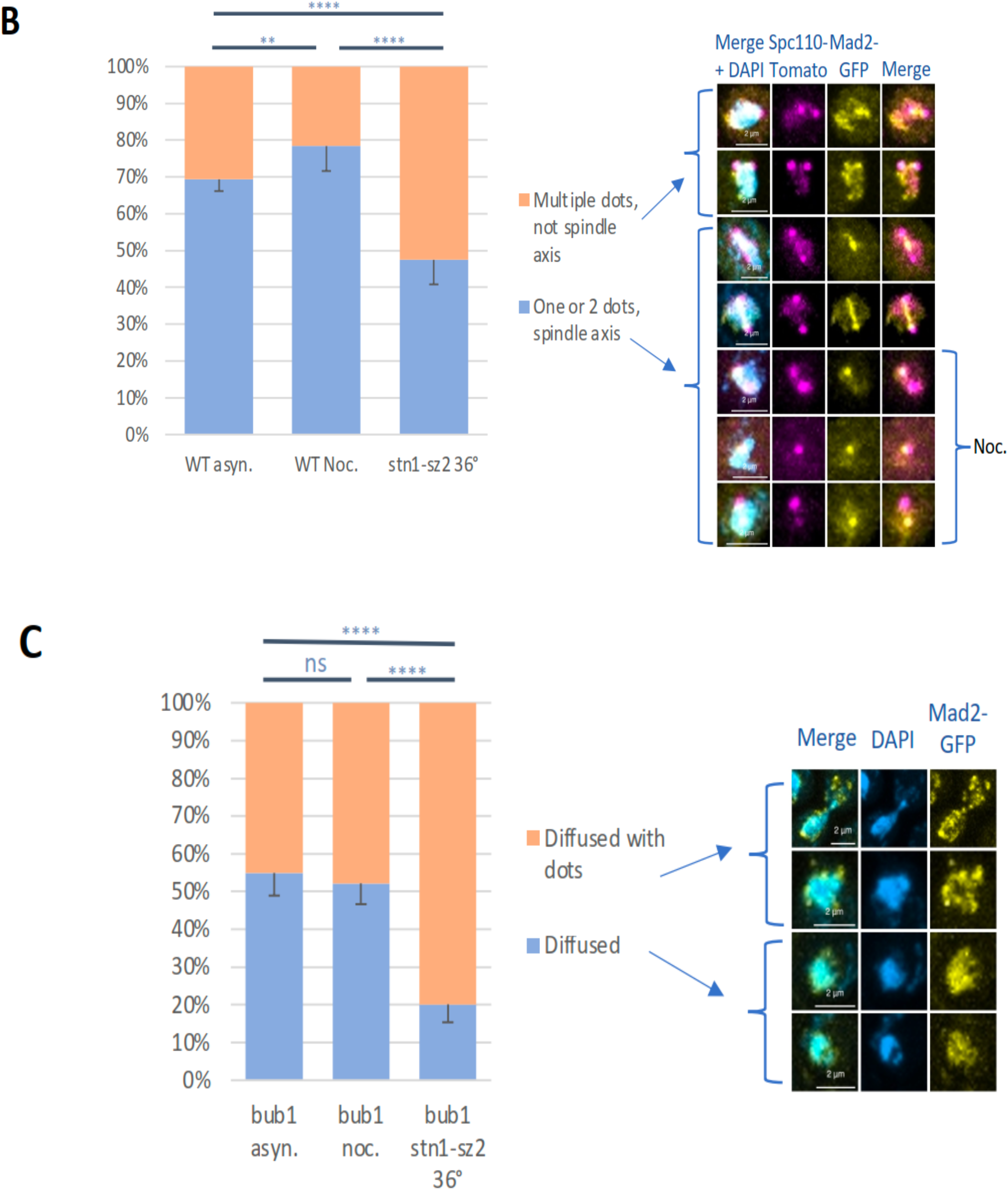
Localization of Mad2-GFP in response to *stn1*-*sz2* or nocodazole spindle damage. Untreated asynchronous wild-type cells (“WT asyn.”) or *BUB1*-deleted cells (“bub1 asyn.”) were used as a control for the temperature-sensitive *stn1*-*sz2* mutant cells grown for 2 hr at the restrictive temperature of 36°C (“stn1-sz2 36°” and “stn1-sz2 bub1 36°”) as well as for the asynchronous wildtype cells treated with 15 μg/ml nocodazole (“WT Noc.” and “bub1 Noc.”). Mad2-GFP (panels A, B and C) and Spc110-Tomato (panel B) constructs were used to detect Mad2 and the SPBs respectively. Pink (instead of red) and yellow (instead of green) were used to show the Tomato and GFP signals, respectively, to allow optimal visualization. The histograms show the quantification of the different patterns of localization of the Mad2-GFP signals in the indicated backgrounds after 2 hr growth at 36°C. (**A**) Results from 12 independent experiments (for “WT asyn.”) and 18 each for “WT Noc.” and “stn1-sz2 36°” are shown as percentages of total cells (100 cells counted per experiment). (**B)** Results from 8 (for “WT asyn.”) and 10 (for “WT Noc.” and “stn1-sz2 36°”) independent experiments are shown as percentages of total cells (100 cells counted per experiment). (**C**) Results from 16 independent experiments for each of the three backgrounds used are shown as percentages of total cells (100 cells counted per experiment). Significance numbers are p-values from the Student’s t-test (**** indicates a p-value < 0.0001).

In wild type, untreated, cells, Mad2-GFP was found to be mostly located in diffuse signals around the nucleus, sometimes with one or more distinct loci with a stronger signal (**Figure 5A**). Strikingly, nocodazole treatment induced a clear re-localization of Mad2-GFP, which was then rather found as concentrated loci, even if some diffuse signals could still be observed around the nucleus (**Figure 5A**). The kinetochores are normally attached to the mitotic spindle, but since nocodazole treatment provokes a complete depolymerization of the spindle, kinetochores remain associated with SPBs (spindle pole bodies, the homologs of centrosomes). Localization of the two SPBs could be achieved following labelling of Spc110-Tomato (**Figure 5B**). Thus, the Mad2-GFP sharp dots seen to re-localize upon nocodazole treatment defined the position of the kinetochores at the SPBs. On the other hand, in *stn1*-*sz2* mutant cells, Mad2-GFP localized as diffuse signals around the nucleus, but also re-localized as loci appearing under the form of sharp dots (**Figure 5A, B**). The few dots observed in wild-type untreated cells were mostly found to coincide with the spindle axis. These probably corresponded to the normal occasional SAC activation during a normal cell cycle, due to minor endogenous damage. In nocodazole-treated wild-type cells also, the large majority of the Mad2 dots co-localized with the SPBs. In *stn1*-*sz2*, less than half of the Mad2-GFP dots localized to the spindle axis (**Figure 5B**), indicating that Mad2 concentrated, partly, at the kinetochores, but also formed loci elsewhere in the nucleus.

The Mad2-GFP dots might correspond or not to the kinetochores. The localization of dots along the spindle axis was an indication that the dots corresponded to kinetochores. However, the kinetochores could be mislocalized in *stn1*-*sz2* cells. We therefore used the fact that, since the kinetochore localization of Mad2 depends on SAC activation, this localization should no longer be visualized if SAC was inactivated. As shown in **Figure 5C**, when SAC was inactivated, using the *bub1* deletion mutant, approximately 50% of the nuclei displayed Mad2-GFP dots in wild type untreated cells, while, similarly, around 50% of nocodazole-treated cells also displayed Mad2-GFP dots. This was in contrast with the situation both in wild type untreated cells with functional SAC that exhibited approximately 20% of Mad2-GFP dots and in nocodazole-treated cells with functional SAC that exhibited around 80% of Mad2-GFP dots (as seen above, Figure 4A). On the other hand, *stn1*-*sz2* cells displayed around 80% of Mad2-GFP dots upon inhibition of SAC by the bub1 mutation, thus highlighting that these 80% of Mad2-GFP dots were SAC-independent. Therefore, there appears to be approximately 30% more *stn1*-*sz2* cells, compared with nocodazole-treated cells, that exhibit SAC-independent Mad2-GFP foci. A hypothesis concerning the origin of these SAC-independent Mad2 foci in the *stn1*-*sz2* mutant is that it activates the so-called metaphase Top2 checkpoint, distinct from the SAC [30].

## 4. Discussion

Telomere loss or dysfunction has dramatic consequences on overall genome stability because the unprotected ends of the chromosomes then become targets for DNA-modifying enzymes. Such a loss of the integrity of the telomere cap, which comprises specialized telomeric proteins arranged around specialized telomeric DNA, can provoke end-to-end fusion and, consequently, chromosome breakage during the next mitotic stage of chromosome segregation. The mutants of telomeric proteins that provoke such phenotypes generate telomeric DNA damage that activates the G2/M DNA damage checkpoint. *S. cerevisiae* yeast telomeres represent a typical example of this, since all the mutants of the major telomere end protection complex, CST (Cdc13-Stn1-Ten1) mainly activate the Mec1/ATR-mediated DNA damage checkpoint [6,7]. However, an exception to this rule was recently found, with the isolation of the *stn1*-*sz2* mutant that had the particularity to principally activate the two major mitotic spindle checkpoints rather than the DNA damage checkpoint [15]. The *stn1*-*sz2* mutant exhibited profound defects in the organization of the mitotic spindle of division and of the kinetochores [15]. The *stn1*-*sz2* mutant therefore represents the prototype of novel mutants of telomeric CST characterized by mitotic-spindle-damage in addition to the classical telomeric-DNA-damage.

### 4.1. Implication of the kinetochore COMA complex in Stn1-controlled telomeric pathways

In the present paper, we pursued our characterization of the phenotypes of the *stn1*-*sz2* mutant [14,15]. We report the identification of an additional actor, namely the kinetochore COMA complex [18,31,32] that appears to sense the *stn1*-*sz2* damage. Thus, in the absence of either Ctf19 or Mcm21 (CENP-P and CENP-O in humans, respectively), *stn1*-*sz2* mutant cells exhibited greatly decreased viability. This could be due to their ability to inappropriately progress through the cell cycle in the presence of telomeric damage. In this hypothesis, intact Ctf19 or Mcm21 would be normally required to limit the consequences of the *stn1*-*sz2* induced-damage. These characteristics are indicative of a role for Ctf19 and Mcm21 in a checkpoint pathway and, interestingly, COMA has been previously implicated in the indirect recruitment of SAC checkpoint proteins [33,34]. Like for all known types of checkpoints, inactivating damage sensing is going to prevent the cell cycle arrest triggered by this damage, which would therefore explain the synthetic growth defects between the *stn1*-*sz2* mutant and the *ctf19* and *mcm21* mutants observed here. Indeed, cells progressing through mitosis with undetected telomeric damage will inevitably undergo death by mitotic catastrophe.

Okp1 forms a complex with Ctf19 and Mcm21 and, since the latter two behave as checkpoints, we propose that the *stn1*-*sz2* damage might neutralize/sequester Okp1 in this complex, thus preventing cell cycle progression, possibly by blocking access to Ndc10 and Ndc80, two additional inner kinetochore components. Upon overexpression of OKP1 in the *stn1*-*sz2* mutant, the resulting increased intracellular Okp1 levels would mimic an absence of *stn1*-*sz2* damage, because it would be as if Okp1 had been freed from the Ctf19-Mcm21-Okp1 complex. Under such conditions, elevated Okp1 levels might inappropriately (because in the presence of *stn1*-*sz2* damage), provoke cell cycle progression which would aggravate the mutant’s growth impairment. Alternatively, increased dosage of Okp1 might re-enforce its binding to Ctf19, thus counteracting the Ctf19-Csm1 binding necessary for anaphase progression (see below), thereby inhibiting growth of the *stn1*-*sz2* mutant.

An additional actor in these mechanisms is represented by Csm1 (**Figure 6**). Csm1 and Lrs4, its partner in the monopolin complex, are essential for rDNA maintenance during metaphase and its segregation during anaphase [22,35,36]. Importantly, Csm1 has been shown to bind Stn1 by two-hybrid, in a genome-wide screen [21], and the *stn1*-*sz2* and *csm1*Δ mutants found to exhibit synthetic growth defects (present data). Interestingly, both the segregation of rDNA and that of telomeric DNA sequences utilize identical pathways, being controlled by FEAR-activated Cdc14 phosphatase, as well as by condensin and cohesin-independent cohesion [37,38]. In addition, Csm1 was shown to bind Ctf19 by two-hybrid, in a genome-wide screen [21]. Most importantly, Csm1-Lrs4 has been shown to co-localize with the Ndc80 kinetochore protein during anaphase only, after being re-located from its nucleolar localization (where the rDNA resides) in metaphase [22]. On this basis, we propose that Csm1 might represent a missing link between Stn1, at the telomeres, and the organization of the kinetochores during mitosis via Ctf19-Mcm21 (**Figure 6A**).

**Figure 6.**
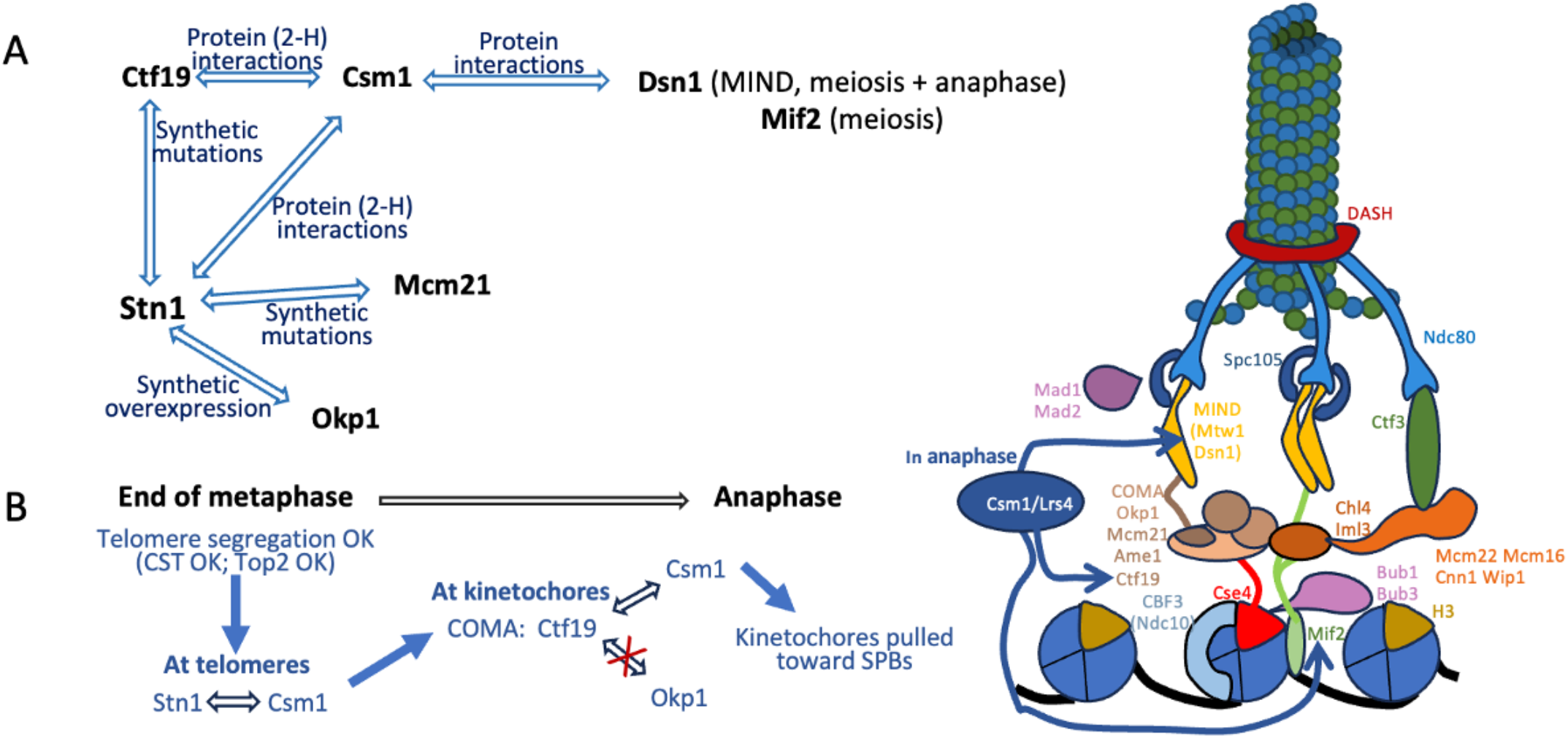
Summary of the interactions between Stn1, COMA, and Csm1. (**A**) Summary of the data presented above and of already published data. (**B**) Working model involving Csm1 and the COMA complex in linking telomere status (damage or not) to kinetochores segregation in anaphase. See text for explanations.

How is Csm1 transiting from the rDNA (and possibly from the telomeres) to the kinetochores is not known at the moment. At this point, an important issue is to understand the role of Csm1 at the kinetochores and how it might be implicated in sensing the *stn1*-*sz2* (or other) damage and act on kinetochore-controlled cell cycle progression. Csm1-Lrs4 has been proposed to act as a molecular clamp, crosslinking kinetochore components to control sister chromatid co-orientation during *S. cerevisiae* meiosis I [39]. Csm1 has loose sequence requirements for binding multiple other proteins, and might therefore explain its implication in, at least, the organization of the nucleolus (where the rDNA localizes), the kinetochores and the Dse3-Mam1-Ulp2-controlled pathways [40]. Potentially, some aspects of telomere maintenance might also be controlled by Csm1, at the same time as it controls rDNA maintenance during metaphase. Incidentally, we note that the fact that Stn1, a telomeric protein, physically binds Csm1 might indicate that Csm1 is needed for the maintenance and segregation of the telomeres as it is for that of the long arm of chromosome XII that contains the rDNA repeats. Both types of repeated sequences have in common to segregate very late during anaphase.

Incidentally, we also note that the rescue of the *stn1*-*sz2* mutant by overexpression of *TOP2* previously observed [14] might be due to Top2 decatenation helping in improving the segregation of the telomeric repeated sequences, as it did for rDNA segregation [38].

A working model is provided in **Figure 6B**. We propose that at the end of metaphase (and maybe earlier), Csm1 binds to telomeric DNA via the CST complex through its Stn1 subunit. In metaphase, in the absence of telomeric damage, sister chromatids are correctly dissociated at telomeric regions. Csm1 protein may then detach from the telomeric regions, as well as, possibly at the same time, detach from the rDNA region, and may then re-localize to the kinetochores **(Figure 6B**). There, Csm1 has been shown to bind Mif2, Dsn1 and Ctf19 in mitosis and/or meiosis [21,39]. We propose that it is the Csm1-Ctf19 interaction that allows to recognize the signal of telomeric status, (“OK for segregation” or “damaged” (not shown in the figure), for instance in *stn1*-*sz2*). This binding of Csm1 to Ctf19 may decrease the binding of Ctf19 to the other units of the COMA complex (Okp1, Mcm21) allowing, by yet unknown mechanisms, the correct kinetochore migration towards the SPBs. This hypothesis would explain how overexpression of *OKP1*, which may counteract the binding of Csm1 to Ctf19, could decrease the amount of Csm1 at the kinetochores, thus inhibiting cell cycle progression in the *stn1*-*sz2* mutant cells. In the *stn1*-*sz2* mutant, telomeres are damaged and therefore their dissociation at the end of metaphase is impaired by checkpoint and recombination/repair mechanisms. The *stn1*-*sz2* mutant protein might sequester Csm1, thus preventing re-localization of the latter to the kinetochores and its binding to Ctf19. This would provoke an abnormal behavior of the kinetochores that would no longer correctly localize to the SPBs during anaphase, as suggested by the defects in the organization of Okp1-GFP observed in the present study.

### 4.2. Stn1-sz2 damage activates a SAC-independent Mad2 checkpoint

In the present study, we have also identified a second novel pathway utilized by the cells in response to *stn1*-*sz2*-generated damage, namely a SAC-independent Mad2-dependent checkpoint. From the literature, the most obvious candidate for this checkpoint might be the so-called Top2 metaphase checkpoint, found to be activated in response to a defect in Top2/TOPOII assembly, or to topoisomerase II inhibitors, both in yeast and humans [30,41]. Other types of defects capable of activating the Top2/TOPOII checkpoint might exist, but have not been identified so far. We suspect that the *stn1*-*sz2* damage might activate the Top2 metaphase checkpoint, due to the previous finding that, like the SAC, it requires Mad2 [30]. In addition, Mad2 must be activated at the kinetochore for the SAC to function, whereas Mad2 was not recruited to kinetochores upon Top2/TOPOII checkpoint activation [30,41]. Likewise, we found here that in the *stn1*-*sz2* mutant less than half of the Mad2 dots localized to the spindle axis, indicating that Mad2, although concentrating, partly, at the kinetochores, also formed loci elsewhere in the nucleus. These data suggested that in the *stn1*-*sz2* mutant, both the SAC and the Mad2-dependent SAC-independent checkpoints were simultaneously activated, as seen by the presence of Mad2-GFP foci both at the kinetochores as well as at places other than at the kinetochores.

Given that Mad2 is known only to participate in two checkpoints, the SAC and the Top2 checkpoint, we propose in **Figure 7** the following model: The *stn1*-*sz2* mutation induces, like all CST mutants, telomeric DNA damage, which prevents the correct segregation of sister chromatids at the end of metaphase, and provokes the recruitment of Top2. However, the topoisomerase fails to properly unwind sister telomeres inducing, in the process, additional damage (e.g. strand breaks). Assuming that the initial DNA damage induced by *stn1*-*sz2* was not sufficient to activate the spindle checkpoint, intervention and failure of Top2 would provide sufficient sister chromatids segregation defect to warn both spindle checkpoints. Mad2, which has an unknown function in the Top2-checkpoint, may be a link between the checkpoint and the SAC in response to some particular damage that Top2 recruitment is unable to repair.

**Figure 7.**
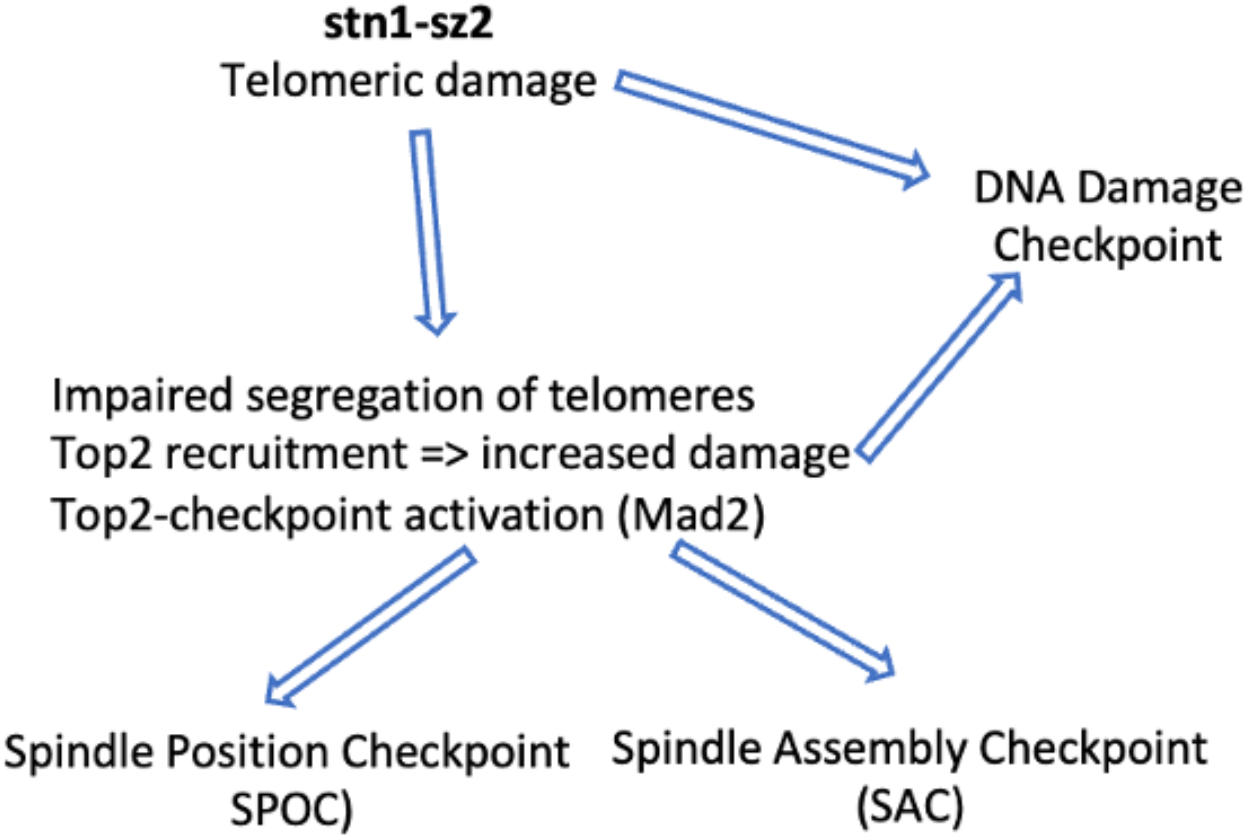
Working model concerning the activation of different checkpoints by *stn1*-*sz2* damage.

In conclusion, the present study allows the opening of new avenues of research on the evolutionary conserved telomeric CST complex, composed of Cdc13/CTC1, Stn1/STN1 and Ten1/TEN1. The major issue of these future studies will be to understand how Stn1 (and potentially Cdc13/CTC1 and TEN1) appears to coordinate the separation of sister chromatids (principally in telomeric regions) with the organization of the mitotic spindle and the kinetochores in order to correctly segregate the previously duplicated genetic material between the mother and daughter cells. It is very likely that the potentially new CST functions described here in budding yeast have been conserved during evolution and also exist in humans [4,42]. In cancer cells, it is particularly important to decipher precise relationships between telomerase reactivation -and maintenance- and the mitotic apparatus. Indeed, both of these deregulated mechanisms have, along the years, become targets of anti-proliferation drugs [4,43].

## Author Contributions

N.G. and M.C. conceived the experiments, acquired funding, conducted the experiments, analyzed the results, and wrote the manuscript.

## Data Availability Statement

All the data supporting the reported results are present in this article. Additional data or protocol details are available upon request.

## Acknowledgments

The authors thank the “LIGUE Nationale contre le Cancer” (CCAURA) for its financial support. We thank Jeffrey Moore, Andrew Murray, Tim Huffaker, and Duncan Clarke for the gifts of strains and plasmids. We also thank the CLIC platform (“Clermont-Ferrand Imagerie Confocale”, Clermont Auvergne University) for help in using the confocal microscopes and associated softwares.

